# Detection of Junctional Ectopic Tachycardia by Central Venous Pressure

**DOI:** 10.1101/2021.04.02.438266

**Authors:** Xin Tan, Yanwan Dai, Ahmed Imtiaz Humayun, Haoze Chen, Genevera I. Allen, Parag N. Jain

## Abstract

Central venous pressure (CVP) is the blood pressure in the venae cavae, near the right atrium of the heart. This signal waveform is commonly collected in clinical settings, and yet there has been limited discussion of using this data for detecting arrhythmia and other cardiac events. In this paper, we develop a signal processing and feature engineering pipeline for CVP waveform analysis. Through a case study on pediatric junctional ectopic tachycardia (JET), we show that our extracted CVP features reliably detect JET with comparable results to the more commonly used electrocardiogram (ECG) features. This machine learning pipeline can thus improve the clinical diagnosis and ICU monitoring of arrhythmia. It also corroborates and complements the ECG-based diagnosis, especially when the ECG measurements are unavailable or corrupted.

## 1 Introduction

Central venous pressure (CVP) is the average blood pressure measured in either superior or inferior vena cava, one of the largest vessels that return blood from the body to the right atrium [13]. This signal reflects the amount of the blood that is returned to the heart [1] and the filling pressure of the right ventricle [5]. The characteristics and amplitude of the CVP waveform components can change significantly with arrhythmias and tricuspid valve pathology [19, 10]. Thus, this signal provides valuable clinical information for arrhythmia diagnosis and automatic detection. However, this signal has been overlooked by most arrhythmia detection literature, which focuses solely on the feature engineering and modeling of ECG signals. This approach can become unreliable when the ECG signal contains artifacts. Thus, a corroborating signal is needed to improve the quality and reliability of an automatic arrhythmia detector.

This paper presents a novel feature extraction and automatic arrhythmia detection pipeline of CVP signal with a focus on pediatric junctional ectopic tachycardia (JET) detection as an exemplar. It contributes to the literature by introducing a CVP signal preprocessing and feature extraction pipeline. We then compare the machine learning model based on extracted CVP features with the one based on gold standard ECG features [20, 18].

The rest of the paper is organized as follows. Section 2 provides a background on the pediatric JET and the clinical basis of JET detection from CVP signal waveforms. Section 3 introduces our method of preprocessing, feature engineering, and machine learning modeling. Section 4 presents the feature performance in both between-subject and within subject-study. In the end, section 5 concludes this paper.

## 2 Background

Congenital heart diseases are among the most common birth defects, affecting ~1% of live births in the United States [3]. Of the postoperative pediatric cardiac patients, up to 48% develop post-operative arrhythmias [12]. JET is one of the most common types of tachyarrhythmia seen during early post-operative care [12] and is very dangerous and difficult to treat in an infant [15]. Currently, there are no automated bedside JET detection methods that are available to clinicians, often leading to delay in diagnosis and subsequent provision of life-saving therapeutic interventions. Most of the current arrhythmia detection algorithms are based on electrocardiogram (ECG) waveforms and result in a staggering number of false alarms (~72-99% of clinical alarms are false [9]). Also, the absence of the P-wave of ECG is considered to be one of the primary morphological features of JET, while current methods cannot robustly detect and measure P-wave, which results in sub-optimal performance for ECG based classifier.

Instead, the JET morphological features are more obvious for the CVP signal. Normal CVP waveforms have 3 systolic components (*c* wave, *x* descent, *v* wave) and 2 diastolic components (*y* descent, *a* wave). During JET, the characteristics and amplitude of the CVP waveform components change significantly. A tall *a* wave, termed a cannon *a* wave is observed [19, 10]. Figure 1 presents a processed and aligned median stack of CVP waveform. Upon on JET onset, the fusion of *a* and *c* wave leads to an obvious cannon *a* wave.

**Fig. 1.**
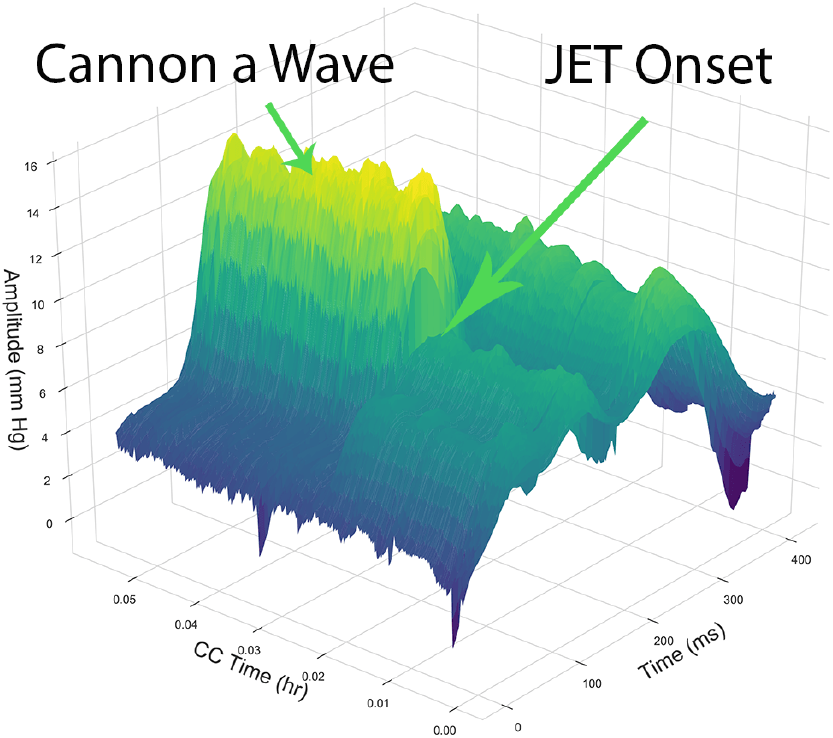
Cannon *a* wave is the primary CVP morphology during JET onset

Figure 2 compares the percentile plot of CVP waveform during JET (left) and sinus waveform (right, sinus refers to normal electrical activity within the heart) with the same scale. 4.5 hours of CVP waveform are used to generate this comparative plot. Significant morphological differences can be observed in the *a* and *c* waveform of the CVP cycle during JET compared to sinus rhythm. Although this morphological difference has been used extensively in clinical diagnosis, there has not been formal methods of extracting these features and build an arrhythmia detection model. The major challenge is that the CVP waveforms are very easily distorted by artifacts occurring through the water-filled, tubing transducer system and by respiration-induced cyclic changes. These artifacts and signal noises make CVP more difficult to analyze than the ECG signal. To solve this issue, we have developed a robust pipeline of removing these artifacts and extract useful features to detect JET onset.

**Fig. 2.**
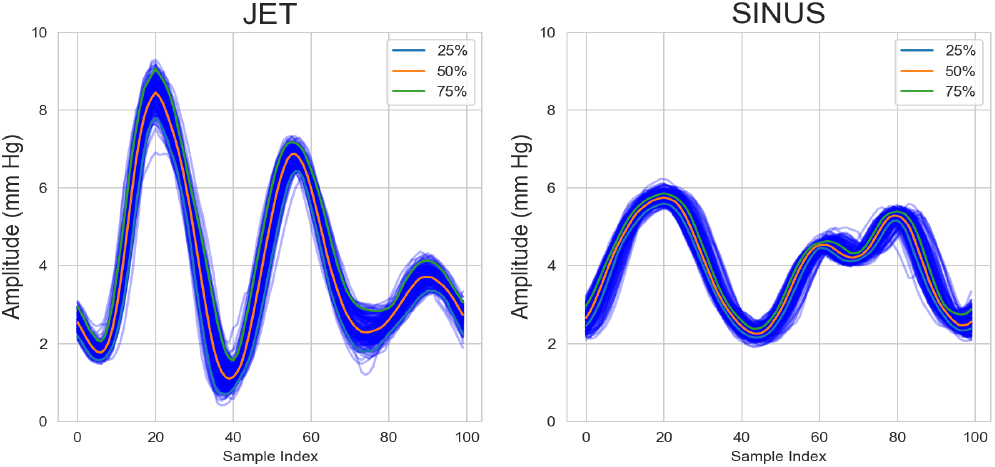
CVP Waveform Comparison

## 3 Methods

In this section, we describe the proposed model for extracting features from CVP signal and detecting JET onset.

### 3.1 Pre-processing

We have developed an elaborate pipeline for CVP data preprocessing, which contains frequency filtering, spike removal, amplitude filter, median filtering, and dynamic alignment. These steps can remove most artifacts and noise in CVP data.

Band filters were modified for CVP data: a 1.5 Hz high pass filter and a 25 Hz low pass filter were used. These two filters can remove the noise caused by infusion, patient breathing patterns, and body movement. Abnormal spikes in segmented 500mm Hg non-overlapping windows were removed using a process introduced by [2]. The raw CVP data is also influenced by multiple noises, which cause the CVP signal to deviate significantly from its theoretical range (2 ~ 6 mmHg). To account for these noises, data points for each patient were ranked in sequence by amplitude, and the values falling outside the 1% − 99% percentile were decreased or increased to this bound.

ECG signals were collected along with the CVP signals. For the ECG signal, WFDB toolbox [21, 11] was used to detect the R-peak and segment each cardiac cycle. Thus, the CVP signal can be segmented and re-sampled accordingly. A median filter with the step of 10 cycles was then applied to reduce the impact of noisy cycles and create a stack. When the ECG signal is unavailable or noisy, the CVP cycle segmentation can still be performed by reference-based waveform matching. Segmenting and stacking the CVP data according to the cardiac cycles results in time offsets because cardiac cycles and CVP cycles are not perfectly synced. In Figure 3 (a), the CVP cycle stacks do not align perfectly. In some scenarios, the offsets can cause peaks in the CVP data to be cut into halves, causing difficulties in feature extraction. Therefore, instead of segmenting and aligning CVP cycles solely by cardiac cycles, ‘dynamic alignment’ is used to align the cycles. This strategy takes the first CVP cycle in the resampled CVP stack as the reference. Then, it realigns every other cycle in the stack according to the position that yields the greatest cross-correlation with the reference stack. In Figure 3 (b), the offsets are removed and the cycles align perfectly with each other. ‘Dynamic alignment’ is superior to the maximum value alignment because it takes into account the entire signal waveform in search for the best position offset.

**Fig. 3.**
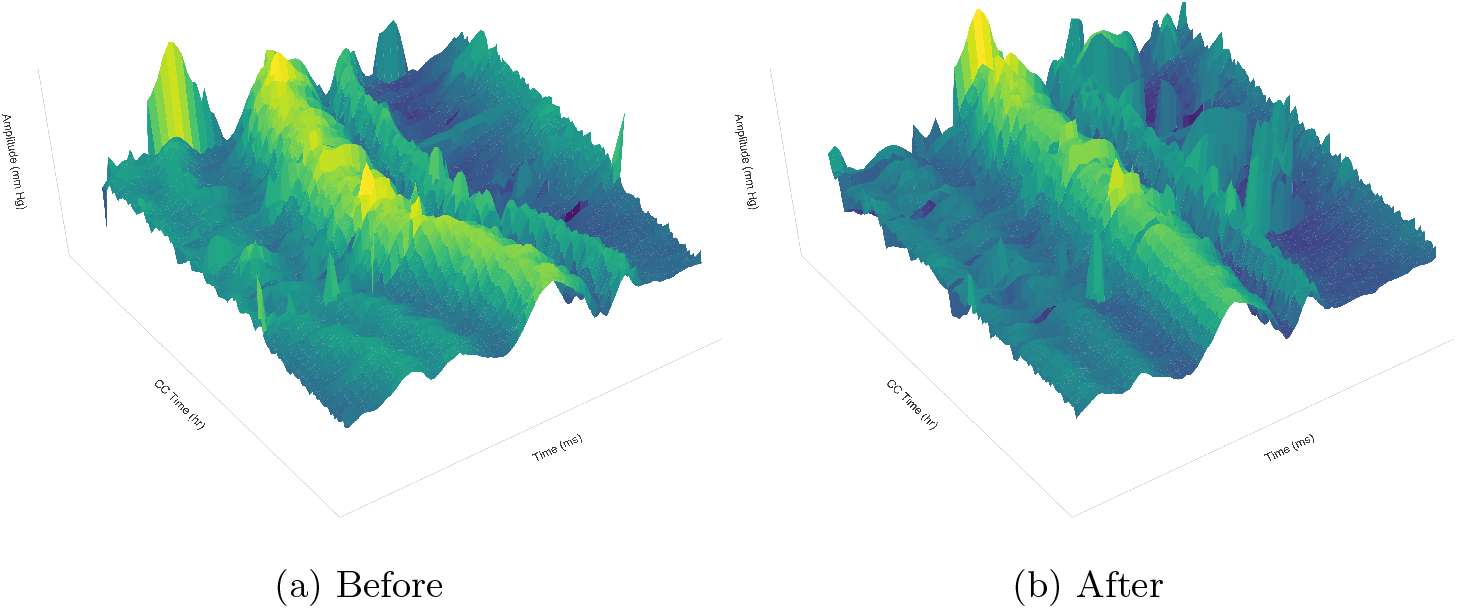
CVP Cycle Dynamic Alignment

### 3.2 Feature Engineering

As demonstrated by clinical study [10, 19] and Figure 1 and 2, the primary morphological feature of JET onset lies in the *a* peak. We propose the following features extraction strategy to measure characteristics of the *a* peak and the overall CVP cycle waveform.

We use four features to characterize the CVP *a* peak. Peak prominence measures how much a peak stands out from the surrounding baseline of the signal. In other words, it is the vertical distance between the peak and its lowest baseline (marked red in Figure 4). The peak height and width are the yellow line and red line identified in Figure 4. The width is identified as the distance between the detected endpoint in the peak, where the height is the distance from the peak to the baseline identified by the endpoint. The width of different relative height can also be obtained. As shown in the graph, the green line represents the peak width at the 50 % level of the peak. The peak slop is calculated as height/width.

**Fig. 4.**
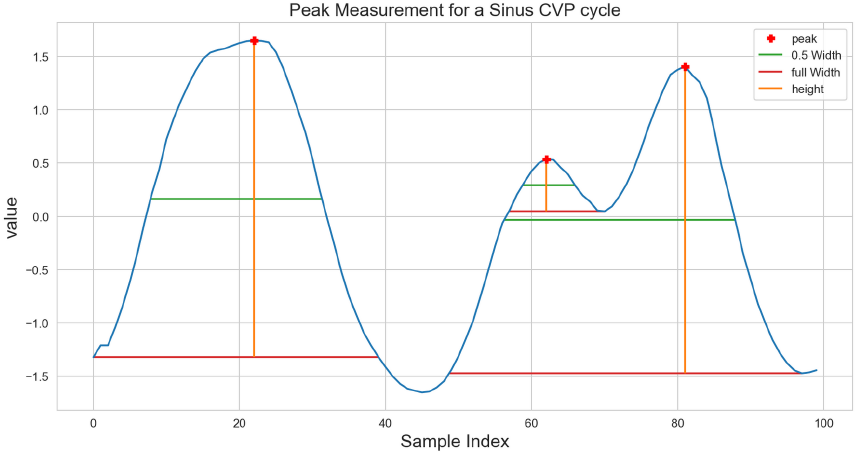
Measure *a, c, v* waves during a single CVP cycle

Beyond features of the *a* wave, we introduce another 5 statistical features to characterize the overall shape of the waveform: waveform mean value, variance, and kurtosis, maximum value, and range. The CVP cycles during JET will display higher mean values and waveform of higher variance [19, 10]. The change in waveform morphology will be reflected in the cycle mean value and variance. Also, since the *a* wave will be disproportionately larger than the other waves in the cycle, the skewness should shift toward the cannon *a* wave. Lastly, we introduce a template-based distance feature, where we measure the cross-correlation between each patient cycle with a standard JET CVP cycle. Despite that each patient has disparate baseline waveform morphology, this distance feature is able to measure the level of similarity between each cycle and a standard typical JET cycle.

These four features can effectively capture the difference between JET and sinus CVP cycles. In Figure 5, feature in each boxplot display clear separation between sinus group and JET group. It demonstrates that the proposed features are able to characterize the morphological difference and provide reliable JET onset detection.

**Fig. 5.**
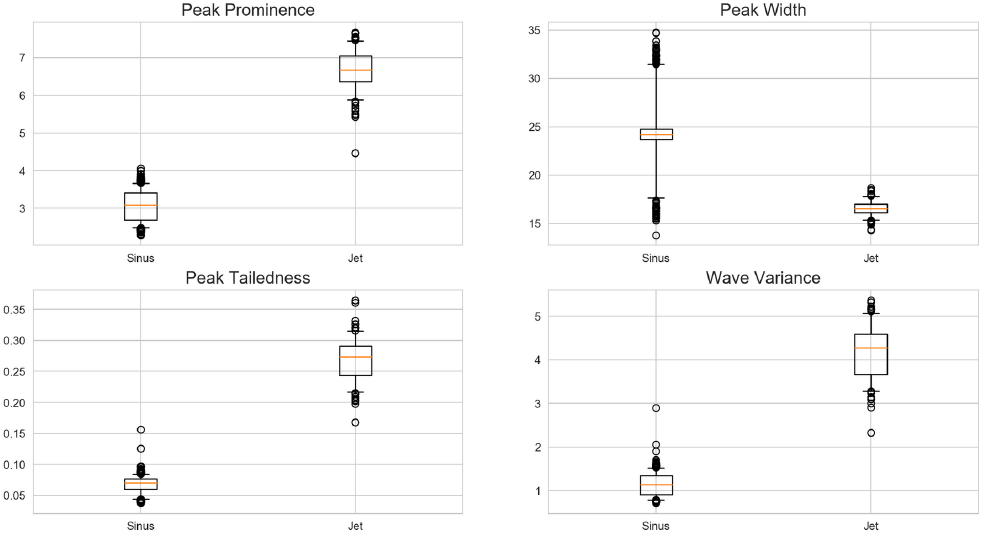
Selected CVP Features Comparison

### 3.3 Models

We use a random forest model [6] with maximum depth of 15 to classify JET vs sinus cardiac cycles from the 10 CVP features and 13 ECG features.

## 4 Experiment and Results

### 4.1 Data Introduction

The data contains 23.3 hours of signal with 6.3 hours of JET and 17 hours of sinus for 8 patients. Throughout the data, there are 4 channels available for the ECG signal and 2 channels for the CVP signal. We only select the channel that contains the best-quality signal to conduct feature engineering and subsequent classification experiment.

### 4.2 Benchmark ECG features

To demonstrate the effect of our proposed features, we compare them against 24 gold-standard ECG features that measure the temporal characteristics of the ventricular depolarization waves (QRS complex) [18, 20]. It includes the following: QRS complex widths, QS width, PR width, Peak Heights (P, R, Q, S), Peak Differences (PQ, RQ, RS), and normalized heart rate features, etc. These are well-validated features characterizing ECG waveform, and they have also been extensively utilized in ECG-based arrhythmia detectors [4, 7, 8, 14, 16, 18, 17].

### 4.3 Experiment Design

We designed two experiments to demonstrate the effectiveness of proposed CVP features. The first experiment conducts within-patient training and testing. The testing data and training data both come from the same patient with a 30%−70% split. The second experiment conducts cross-patient training and testing. In this experiment, the testing data comes from a single patient, and the training data comes from every other patient in the dataset. For each experiment, we report the sensitivity, specificity, and area under curve (AUC) of the random forest model trained with CVP features, ECG features, and CVP + ECG features combined respectively. Finally, we also report the average feature importance of CVP and ECG features in the joint model. Thus, we can compare the importance of proposed CVP features versus ECG features.

### 4.4 Results

In both experiments, the model relying on CVP features alone achieves comparable performance with the model relying on ECG features. The within-patient experiment generally yields better performance than the cross-patient experiment. The reason is that each patient has underlying diseases, which creates a morphological disparity in the CVP waveform. Despite the waveform disparity, the performance of the model relying on CVP features still matched the performance of the model relying on ECG features. As shown in Figure 6, when ECG features and CVP features are jointly utilized in the model, CVP features have importance scores than the ECG features.

**Fig. 6.**
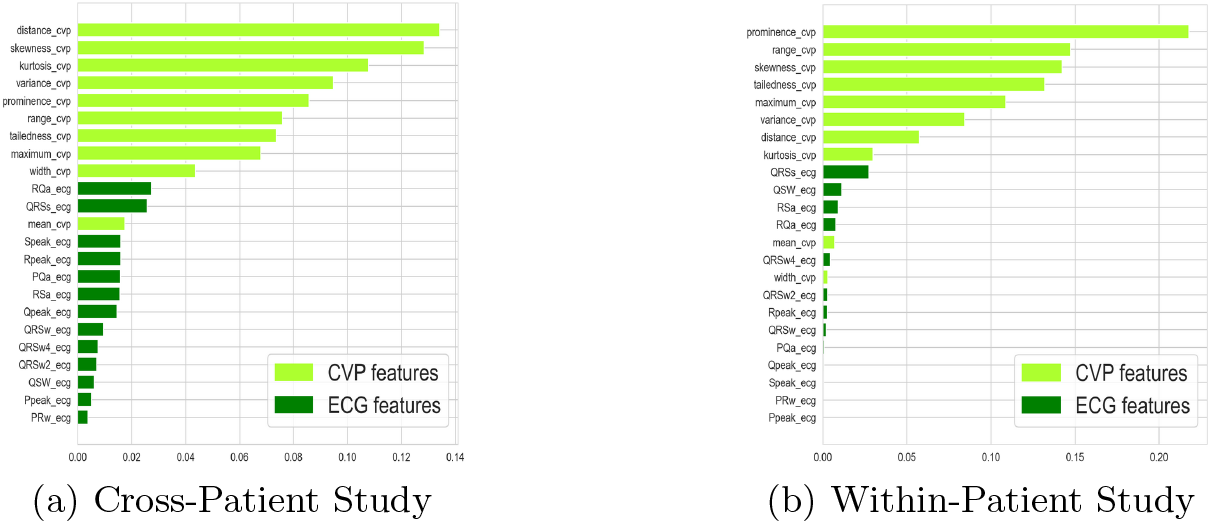
Feature Importance Scores

### 4.5 Limitations

As mentioned in the introduction, changes in CVP could be due to triscuspid pathology. However, the patients in this analysis all had JET as the inciting event for changes in CVP. Additionally, we looked at relative change in CVP morphology due to JET, which takes into account any pre-existing tricuspid valve abnormalities. Usually, more than one channel of CVP signal will be available. However, this model only selects the single signal channel with the best quality to experiment. Future work can focus on developing models for multichannel signals.

**Table 1.** Within-Patient Experiment Result

| Test Patient: | 1 | 2 | 3 | 4 | 5 | 6 | 7 | 8 |
| --- | --- | --- | --- | --- | --- | --- | --- | --- |
| CVP features |  |  |  |  |  |  |  |  |
| Sensitivity | 1 | 0.99 | 1 | 1 | 0.53 | 1 | 0.95 | 1 |
| Specificity | 1 | 0.91 | 0.98 | 1 | 0.99 | 1 | 1 | 1 |
| AUC | 1 | 0.99 | 0.99 | 1 | 0.94 | 0.99 | 0.98 | 1 |
| ECG features |  |  |  |  |  |  |  |  |
| Sensitivity | 1 | 1 | 1 | 1 | 0.79 | 0.98 | 0.98 | 1 |
| Specificity | 1 | 1 | 1 | 1 | 1 | 1 | 1 | 1 |
| AUC | 1 | 1 | 1 | 1 | 0.99 | 0.99 | 1 | 1 |
| ECG + CVP |  |  |  |  |  |  |  |  |
| Sensitivity | 1 | 1 | 0.98 | 1 | 0.86 | 0.98 | 0.94 | 0.99 |
| Specificity | 1 | 1 | 1 | 1 | 1 | 1 | 1 | 1 |
| AUC | 1 | 1 | 1 | 1 | 0.98 | 0.99 | 0.99 | 1 |

**Table 2.**
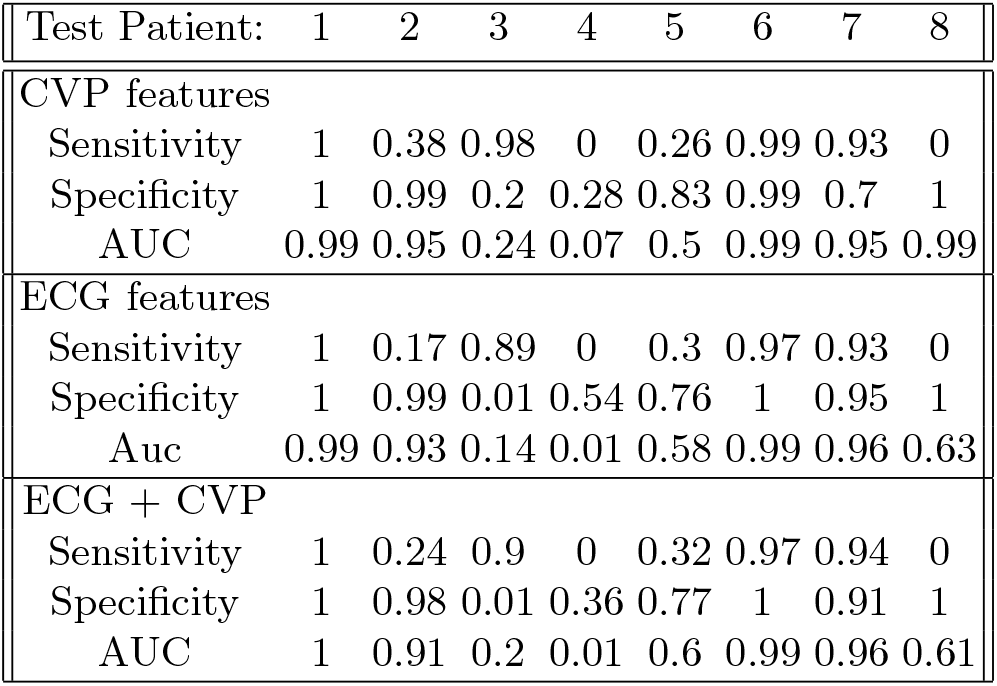
Cross-Patient Experiment Result

| Test Patient: | 1 | 2 | 3 | 4 | 5 | 6 | 7 | 8 |
| --- | --- | --- | --- | --- | --- | --- | --- | --- |
| CVP features |  |  |  |  |  |  |  |  |
| Sensitivity | 1 | 0.38 | 0.98 | 0 | 0.26 | 0.99 | 0.93 | 0 |
| Specificity | 1 | 0.99 | 0.2 | 0.28 | 0.83 | 0.99 | 0.7 | 1 |
| AUC | 0.99 | 0.95 | 0.24 | 0.07 | 0.5 | 0.99 | 0.95 | 0.99 |
| ECG features |  |  |  |  |  |  |  |  |
| Sensitivity | 1 | 0.17 | 0.89 | 0 | 0.3 | 0.97 | 0.93 | 0 |
| Specificity | 1 | 0.99 | 0.01 | 0.54 | 0.76 | 1 | 0.95 | 1 |
| Auc | 0.99 | 0.93 | 0.14 | 0.01 | 0.58 | 0.99 | 0.96 | 0.63 |
| ECG + CVP |  |  |  |  |  |  |  |  |
| Sensitivity | 1 | 0.24 | 0.9 | 0 | 0.32 | 0.97 | 0.94 | 0 |
| Specificity | 1 | 0.98 | 0.01 | 0.36 | 0.77 | 1 | 0.91 | 1 |
| AUC | 1 | 0.91 | 0.2 | 0.01 | 0.6 | 0.99 | 0.96 | 0.61 |

Morphological disparity across different patients exist in both ECG and CVP signal. Despite that the JET vs sinus cycles are easily separable during the within-patient experiment, the morphological disparity undermines the cross-patient experiment result. The reason could be that the patients have other cardiac disease or they are under certain medication, which complicate the waveform.

## 5 Conclusion

We have proposed a novel pipeline to process and extract features from Central Venous Pressure with a case study on the automatic detection of junctional ectopic tachycardia. The preprocessing pipeline and feature engineering pipeline provide a solution to remove complex artifacts in the CVP waveform and extract clinically valuable information. The within-patient and cross-patient experiment demonstrate that the CVP signal is as reliable as the ECG signal in detecting JET onset, and CVP features have higher importance scores than ECG features in the joint model. Thus, the quality and performance of arrhythmia detectors can be improved by incorporating CVP signals, and it can be particularly important when ECG signals are unavailable or contain major artifacts.

## 6 Acknowledgements

Genevera Allen acknowledges support from NSF DMS-1554821, NSF NeuroNex-1707400, and NIH 1R01GM140468.

## References

1. Central venous pressure – evaluation, interpretation, monitoring, clinical implications. Bratisl Lek Listy 109(4), 185–187

2. Segmentation of heart sound recordings by a duration-dependent hidden markov model. Physiological Measurement 31(4), 513–529 (2011)

3. Data and statistics on congenital heart defects — cdc (Nov 2018), https://www.cdc.gov/ncbddd/heartdefects/data.html

4. Abawajy, J., Kelarev, A., Chowdhury, M.: Multistage approach for clustering and classification of ecg data. Computer Methods and Programs in Biomedicine 112, 720–730 (12 2013). https://doi.org/10.1016/j.cmpb.2013.08.002

5. Bigatello, L.M., George, E.: Hemodynamic monitoring. Minerva Anestesiologica 68(4), 219–225 (Apr 2002), https://pubmed.ncbi.nlm.nih.gov/12024086/

6. Breiman, L.: Random forests. Machine Learning 45(1), 5–32 (2001). https://doi.org/10.1023/a:1010933404324

7. Chang, C.C., Lin, C.J.: Libsvm. ACM Transactions on Intelligent Systems and Technology 2, 1–27 (04 2011). https://doi.org/10.1145/1961189.1961199

8. deChazal, P., O’Dwyer, M., Reilly, R.: Automatic classification of heartbeats using ecg morphology and heartbeat interval features. IEEE Transactions on Biomedical Engineering 51, 1196–1206 (07 2004). https://doi.org/10.1109/tbme.2004.827359

9. Drew, B.J., Harris, P., Zègre-Hemsey, J.K., Mammone, T., Schindler, D., Salas-Boni, R., Bai, Y., Tinoco, A., Ding, Q., Hu, X.: Insights into the problem of alarm fatigue with physiologic monitor devices: A comprehensive observational study of consecutive intensive care unit patients. PLoS ONE 9(10), e110274 (Oct 2014). https://doi.org/10.1371/journal.pone.0110274

10. Fujita, Y., Hayashi, D., Wada, S., Yoshioka, N., Yasukawa, T., Pestel, G.: Central venous pulse pressure analysis using an r-synchronized pressure measurement system. Journal of Clinical Monitoring and Computing 20(6), 385 (Oct 2006). https://doi.org/10.1007/s10877-006-9035-y

11. Goldberger, A., Amaral, L., Glass, L., Hausdorff, J., Ivanov, P., Mark, R., Mietus, J., Moody, G., Peng, C., Stanley, H.: Physiobank, physiotoolkit, and physionet: components of a new research resource for complex physiologic signals. Circulation 101(23), E215—20 (June 2000). https://doi.org/10.1161/01.cir.101.23.e215, https://doi.org/10.1161/01.cir.101.23.e215

12. Kabbani, M.S., Taweel, H.A., Kabbani, N., Ghamdi, S.A.: Critical arrythmia in postoperative cardiac children: Recognition and management. Avicenna journal of medicine 7(3), 88–95 (Jul 2017)

13. Klabunde, R.K.: Central venous pressure. Image for Cardiovascular Physiology Concepts (Apr 2014), https://www.cvphysiology.com/BloodPressure/BP020

14. Korürek, M., Doğan, B.: Ecg beat classification using particle swarm optimization and radial basis function neural network. Expert Systems with Applications 37, 7563–7569 (12 2010). https://doi.org/10.1016/j.eswa.2010.04.087

15. Kylat, R.I., Samson, R.A.: Junctional ectopic tachycardia in infants and children. Journal of Arrhythmia 36(1), 59–66 (Dec 2019). https://doi.org/10.1002/joa3.12282

16. Llamedo, M., Martínez, J.P.: Heartbeat classification using feature selection driven by database generalization criteria. IEEE Transactions on Biomedical Engineering 58, 616–625 (03 2011). https://doi.org/10.1109/tbme.2010.2068048

17. Luz, E.J.D.S., Schwartz, W.R., Cámara-Chávez, G., Menotti, D.: Ecg-based heartbeat classification for arrhythmia detection: A survey. Computer Methods and Programs in Biomedicine 127, 144–164 (2016). https://doi.org/10.1016/j.cmpb.2015.12.008

18. Mar, T., Zaunseder, S., Martinez, J.P., Llamedo, M., Poll, R.: Optimization of ecg classification by means of feature selection. IEEE Transactions on Biomedical Engineering 58, 2168–2177 (08 2011). https://doi.org/10.1109/tbme.2011.2113395

19. Pittman, J.A.L., Ping, J.S., Mark, J.B.: Arterial and central venous pressure monitoring. International Anesthesiology Clinics 42, 13–30 (2004). https://doi.org/10.1097/00004311-200404210-00004

20. Saenz-Cogollo, J.F., Agelli, M.: Investigating feature selection and random forests for inter-patient heartbeat classification. Algorithms 13, 75 (04 2020). https://doi.org/10.3390/a13040075, https://www.mdpi.com/1999-4893/13/4/75

21. Silva, I., Moody, G.B.: An open-source toolbox for analysing and processing physionet databases in matlab and octave. Journal of Open Research Software 2 (09 2014). https://doi.org/10.5334/jors.bi

